# Transcranial Random Aperiodic Stimulation Improves Working Memory Precision

**DOI:** 10.64898/2026.09.08.750163

**Authors:** Quirine van Engen, Justin Riddle, Bradley Voytek

**Affiliations:** Department of Cognitive Science, University of California, San Diego; Department of Psychology, Florida State University; Program in Neuroscience, Florida State University; Halıcıoğlu Data Science Institute, University of California, San Diego; Neurosciences Graduate Program, University of California, San Diego; Kavli institute for Brain and Mind, University of California, San Diego

**Author notes:** These authors contributed equally to this work (Justin Riddle, Bradley Voytek).

## Abstract

Aperiodic neural activity – historically dismissed as noise – is independently modulated from neural oscillations during visual working memory maintenance. In the context of aging, “flatter” trait aperiodic activity is associated with poorer visual working memory performance. Here, we causally test the role of aperiodic neural activity in visual working memory in younger adults. We introduce a novel noninvasive neurostimulation method – transcranial random aperiodic stimulation (tRAS) – to causally manipulate aperiodic activity to be either steeper or flatter. In a randomized, double-blind, placebo-controlled, crossover neurostimulation design (n = 30), we show that online noninvasive tRAS improves visual working memory precision when aperiodic activity is causally steepened. Our novel stimulation method paves the way for causal studies of non-oscillatory, aperiodic activity in human cognition, aging, and disease.

## Main Text

When planning your next move in a board game, you need to remember the various context-dependent rules while simulating multiple moves into the future. A key cognitive function supporting goal-directed behavior is working memory (WM) – the ability to maintain information that is no longer in perception (*1, 2*). Critically, WM capacity is limited (*3–5*). For example, remembering the positions of eight pieces in a board game is much more difficult than three.

Many WM studies leverage noninvasive electroencephalography (EEG) to investigate network-level electrical activity correlated with WM processes (*6*). Such studies have identified frontal-midline theta activity (4-8 Hz) as a robust correlate of WM encoding, maintenance, and retrieval (*7–11*). Beyond these correlational analyses, causal manipulation of theta-frequency network activity using noninvasive electrical stimulation has selectively increased or decreased coupling between lateral prefrontal and posterior parietal cortex and found a corresponding increase or decrease in behavioral indices of WM (*12–15*).

However, theta oscillations are surprisingly scarce, only manifesting on a subset of trials and does not exceed background levels of activity in the resting-state for a majority of participants (*16–19*). Notably, neural oscillations are embedded in an electrophysiological background, characterized as aperiodic activity and manifesting as unstructured voltage fluctuations over time. Aperiodic activity is reflected in the frequency domain as a downward slope, where low-frequency activity has greater power than high-frequency activity (Fig. 1A). While aperiodic activity was historically considered noise, studies investigating task-based aperiodic shifts confirm that it systematically changes during visual WM (*17, 20, 21*). These aperiodic activity shifts are distinct from theta power shifts (*22*), with evidence for a task-general decrease in aperiodic activity regardless of WM load (*23*). Methodological limitations of narrow-band filtering techniques could mistake a steepening of aperiodic activity as an increase in amplitude of theta oscillations, even though no oscillations are present in the signal (Fig. 1B). We investigated whether non-invasive electrical stimulation targeting aperiodic activity in the lateral fronto-parietal network (Fig. 1C&D) would alter WM precision on a continuous response delayed-recall WM task (Fig. 1F). Utilizing a two-session crossover experimental design, baseline WM capacity was estimated during high-density EEG. Then, in the second session, participants received a novel form of aperiodic neurostimulation, transcranial random aperiodic stimulation (tRAS), while performing a titrated version of the visual WM task as EEG was acquired (Fig. 1 E&F). With steep-tRAS, random electrical noise was delivered with a steeper than endogenous 1/f noise distribution, whereas flat-tRAS utilized a flatter slope that was closer to white noise. We predicted that steep-tRAS relative to placebo would improve behavioral indices of WM and would increase the slope of the aperiodic slope measured during interleaved resting-state recordings.

**Fig. 1:**
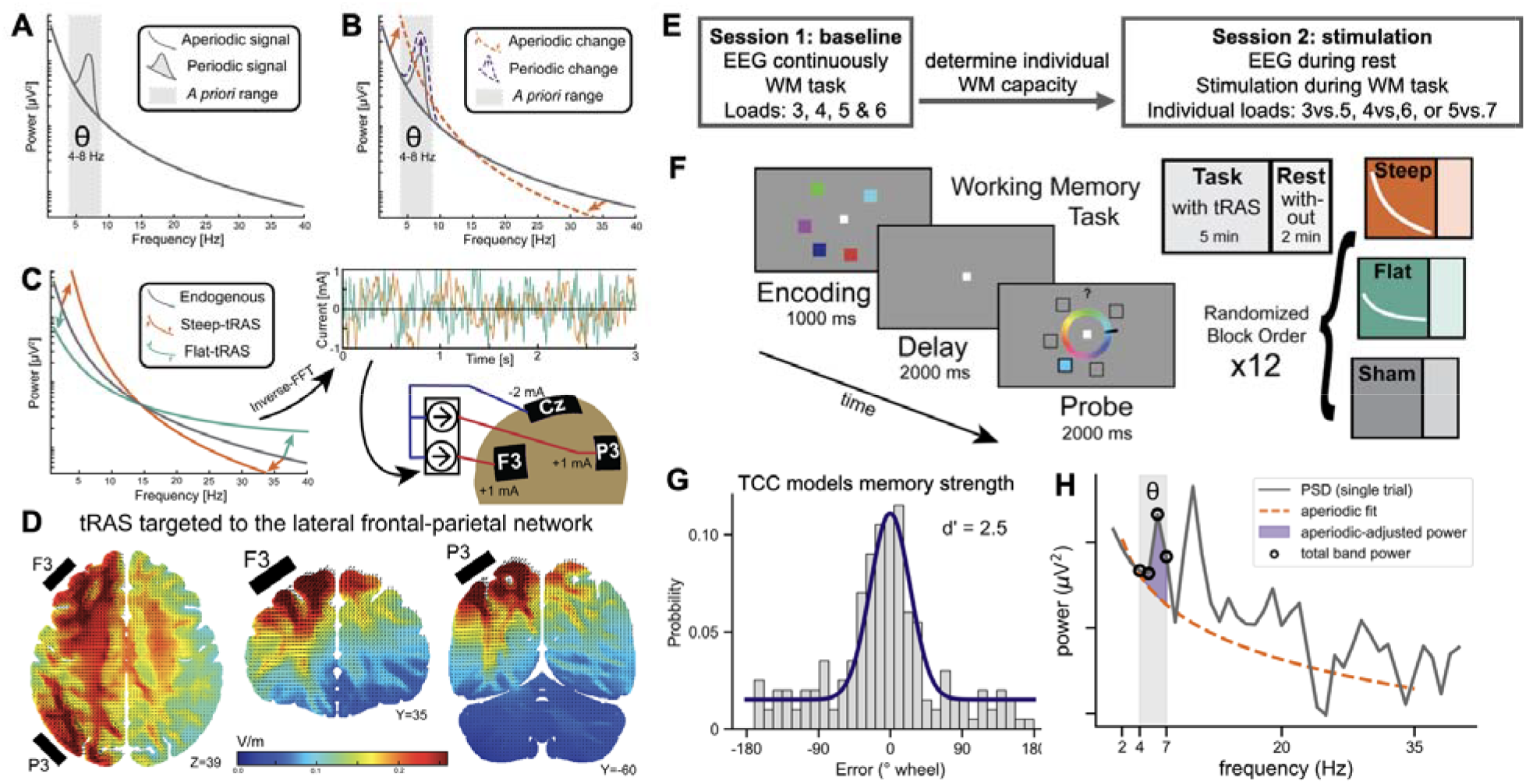
Study overview. (A) Schematic power spectral density plot (PSD) where periodic theta power (oscillatory) manifests as a “bump” on top of the aperiodic signal. (B) Hypothesized methodological measurement error in which an increase in theta bandpower could be either driven by an increase in aperiodic-corrected theta power (reflecting putative oscillations), or an increase in power due to aperiodic steepening. (C) Approach for creating novel aperiodic stimulation method. The left depicts aperiodic activity in the frequency domain, in which the exponent of the flat-tRAS is chosen to be flatter than the average endogenous activity, and steep-tRAS was steeper than average. The inverse-FFT transformed these signals to the time domain, and these were used as noninvasive electrical stimulation waveforms. (D) Based on the electrode montage setup, modeled electrical field potentials suggest that the highest electric field strength was in the fronto-parietal network. (E) Overview of the experimental design. Session 1 served as a baseline measurement to determine individual WM capacity and task difficulty for session 2. Participants receive stimulation during the WM task in session 2, and EEG is measured during the rest periods. (F) Continuous response, delayed recall WM task. This task was used in both sessions and was titrated for the second session. Either the standard 3, 4, 5 or 6 items, or individualized low and high loads. In session 2, participants received stimulation while performing the task in a randomized order of either steep-tRAS, flat-tRAS, or sham. (G) WM precision error was calculated as the difference between the probed and reported color in degrees, where d′ represents memory precision. A higher d′ indicates better memory precision. (H) The aperiodic signal was quantified from pre-stimulus and WM delay periods in session 1 using the spectral parameterization toolbox (*24*).

## Results

### WM precision decreased with greater memory load

Thirty-five participants completed the baseline session, from which 31 were included in the behavioral analysis. Behavioral performance was analyzed using a linear mixed model (LMM). Response time was significantly increased with WM load (β = -0.009, p = 4.4*10^-3^, 95%CI = [-0.003, -0.015]) (Fig. S1A). We estimated WM precision (d’) using the target confusability competition (TCC) model on the distribution of degrees of distance between the probed and reported color (Fig. 1H). TCC accounts for perceptual similarity between stimuli (*25*).. We found that WM precision decreased as a function of WM load (3, 4, 5, or 6 items) (β = -0.47, p = 2.00*10^-38^, 95%CI = [-0.52, -0.43], Fig. 2A).

**Fig. 2:**
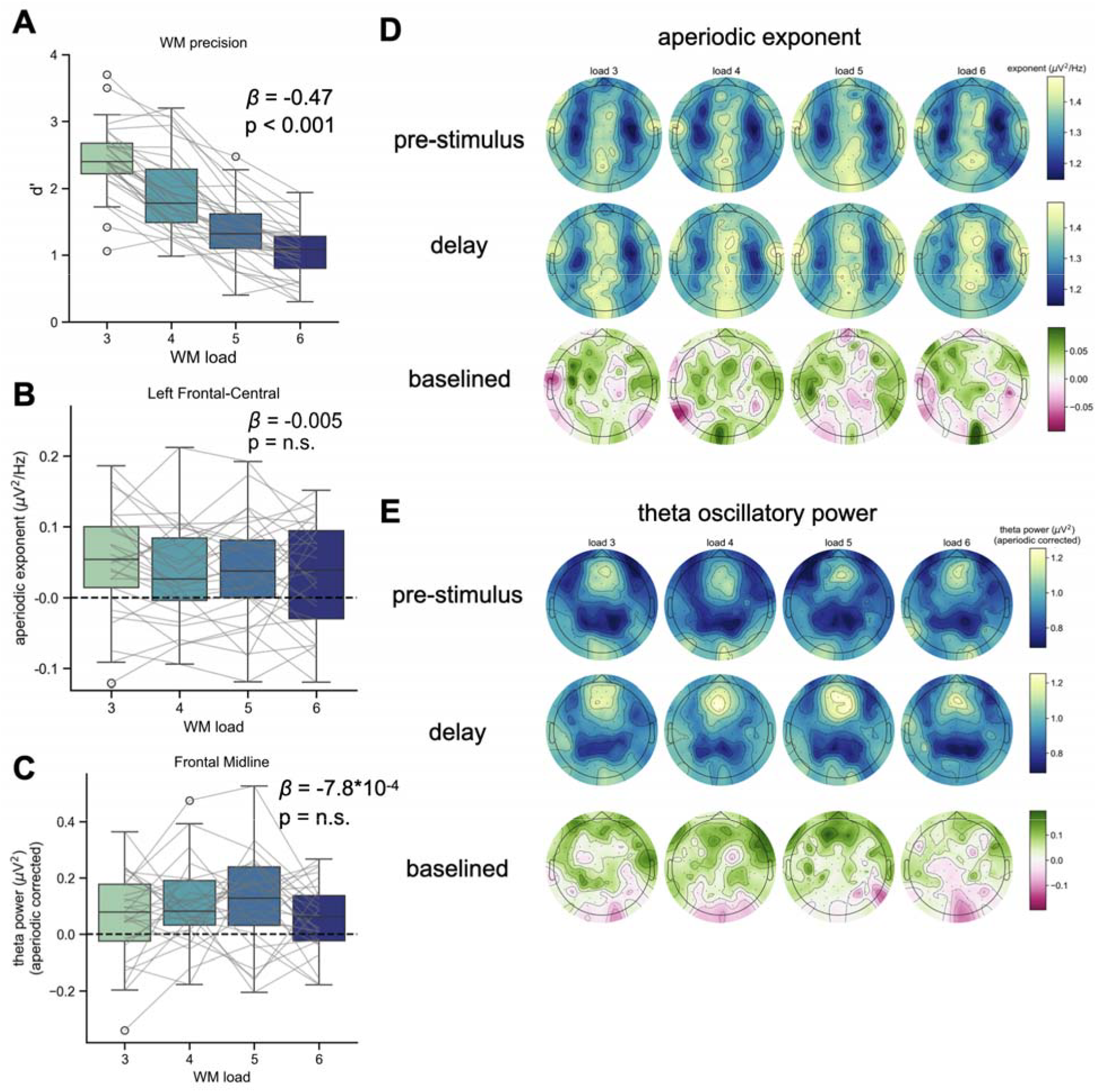
Baseline session, behavioral and EEG results. (A) WM precision significantly decreased as WM load increased. (B) Fronto-central aperiodic activity was systematically steeper during the delay period compared to baseline but did not change with WM load. (C) Aperiodic-adjusted frontal-midline theta power was unaffected by WM load. (D) Topographic analysis for the aperiodic exponent is shown for the pre-stimulus, delay period, and the baselined (delay minus pre-stimulus) period revealed a pattern of steeper aperiodic activity over the midline, and flatter bilaterally over all loads. (E) Topographic analysis of the aperiodic-corrected theta power revealed a clear frontal-midline cluster, but activity did not change with WM load.

### Aperiodic activity and frontal-midline theta oscillations were unaffected by WM load

From the 31 participants, two were further excluded from the EEG analysis due to poor EEG quality, resulting in 29 participants. To investigate WM load dependent changes in aperiodic activity and power of frontal-midline theta oscillations, we applied spectral parameterization during the pre-stimulus and delay periods of the WM task. Topographic analysis of the aperiodic exponent (Fig. 2D) indicated that a frontal-central region was modulated by the task. Thus, aperiodic activity was further investigated over the frontal-central electrode cluster. The model including load as a fixed factor was not significantly different from the model with only random intercepts to account for between-subject variability. WM load did not affect the aperiodic exponent during the WM delay period (β = -0.005, p = 0.16, 95%CI = [-0.012, 0.002]), but the aperiodic exponent was significantly increased (steeper) during the delay period from pre-stimulus as determined by the intercept of this model (intercept = 0.063, p = 2.5*10^-3^, 95%CI = [0.023, 0.102]) (Fig. 2B). The increase in delay-period aperiodic exponent in frontal electrodes is consistent with a previous study using a similar delay-recall task (*23*), and suggests that stimulation designed to increase the aperiodic exponent should facilitate WM-related task performance.

Frontal-midline theta activity was analyzed over frontal-midline electrodes, correcting for the influence of aperiodic activity to isolate putative theta oscillations. The topoplots during the baseline and delay period show a stereotypical theta power distribution that is focused on the frontal-midline (Fig. 2E). The model for aperiodic-corrected theta power did not show a significant effect of WM load (β = -7.8*10^-4^, p = 0.93, 95%CI = [-0.02, 0.019]) (Fig. 2C). These results are consistent with an emerging literature regarding the potential confound of aperiodic activity on the interpretation of power changes in theta oscillations during visual WM (*17, 22*).

Previous work has also discovered an association between resting-state aperiodic activity and WM performance (*26–28*), we investigated whether trait-level aperiodic exponent was predictive of individual difference in WM precision. We computed the aperiodic exponent during the eyes-open resting periods in between blocks and correlated this with the average WM precision over all WM loads. No evidence was found that aperiodic activity at rest was associated with overall WM precision (r(29) = -0.22, p = 0.26, 95%CI = [-0.54, 0.16]) (Fig. S2).

### Steeper-tRAS improves WM precision

Based on the baseline session, visual WM loads were titrated for each individual. Thirty participants completed the stimulation session where aperiodic neurostimulation, tRAS, was delivered while participants performed the WM task. Fixed values were used for the aperiodic exponent of tRAS. Investigation of the baseline aperiodic exponent confirmed that tRAS was steeper and flatter than the aperiodic exponent of each participant with only a couple exceptions (Fig. S3). After each WM task block, a resting-state period was recorded to investigate the lasting impact of tRAS. Twenty-seven participants were included in the behavioral analysis (2 excluded for at chance task performance, and one did not complete the session) and 26 were included in the EEG resting-state analysis (1 additional excluded for poor EEG quality).

Like the baseline session, we observed a significant main effect of WM load on WM precision (low = 1.97±0.64, high = 1.11±0.36, F(1, 26) = 118, p = 3.5*10^-11^, η _p_ ^2^ = 0.82). Critically, there was also a significant main effect of stimulation type on WM precision (flat-tRAS = 1.50±0.67, sham = 1.52±0.66, steep-tRAS = 1.60±0.71, F(2, 52) = 3.70, p = 0.031, η_p_^2^ = 0.12) (Fig. 3A). Post-hoc tests showed trends toward greater WM precision for steep-tRAS relative to flat-tRAS (t(26) = -2.51, p = 0.056, Cohen’s d = -0.20) and sham-tRAS (t(26) = -2.14, p = 0.084, Cohen’s d = -0.19). Analysis of the difference in WM precision for steep-tRAS relative to flat-tRAS revealed that two thirds of the participants show the hypothesized effect of aperiodic stimulation for both high and low WM loads (Fig. 3B).

**Fig. 3:**
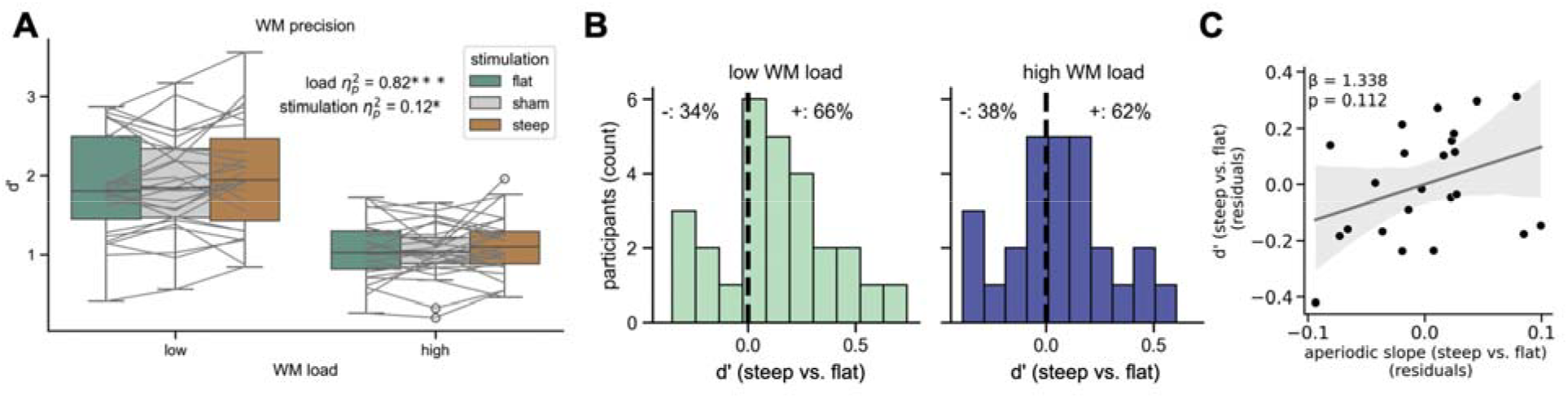
Steep-tRAS improved WM precision compared to sham- and flat-tRAS. (A) WM precision was significantly lower in the high WM-, compared to the low WM-load conditions. Steep-tRAS significantly increased WM precision compared to sham- and flat-tRAS. (B) For both the low and high WM loads, approximately two thirds of the participants showed an increase in WM precision for the steep-versus flat-tRAS conditions. (C) The effect of tRAS on WM precision and aperiodic exponent showed a positive, yet not significant, correlation.

### tRAS did not produce a lasting impact on the aperiodic exponent

To test whether tRAS had a lasting effect on post-stimulation EEG, we quantified aperiodic activity during the 2-minutes of eyes-open resting-state each block after tRAS was completed. Because we hypothesized that tRAS effects would be strongest immediately following stimulation, we investigated the first 30 seconds, as well as the full 2-minutes, of resting state EEG. We did not find any significant differences in aperiodic exponent after stimulation between steep-tRAS, flat-tRAS, and sham-tRAS for either the first 30 seconds after stimulation, nor the full resting period, over all electrodes, the frontal-central cluster, nor the frontal-midline cluster (F < 1) (Fig. S4), suggesting that our tRAS stimulation protocol did not produce a lasting impact on the aperiodic exponent outside of the stimulation window.

### Neither the aperiodic exponent at rest, nor the change due to tRAS predicted WM precision

Lastly, we examined whether WM precision could be predicted from the aperiodic exponent at rest, or from the change in aperiodic exponent between active stimulation conditions. We hypothesized that participants with naturally steeper aperiodic activity at rest would respond differently to tRAS, using an ordinary least squares regression with the aperiodic exponent at rest from the baseline session, and the difference in exponent between the steep-tRAS and flat-tRAS condition. The model was not significant (R^2^ _adj_ = 0.033, p = 0.275). A steepened exponent due to tRAS was positively, but not significantly, related to an increase in WM precision (β = 1.34, p = 0.11, 95%CI = [-0.34, 3.02])(Fig. 3C). The aperiodic exponent at rest from the baseline session did not predict the changes in WM precision induced by tRAS (β = -0.09, p = 0.72, 95%CI = [-0.58, 0.41]).

## Discussion

We introduced a novel, non-invasive aperiodic neurostimulation protocol – tRAS – to test if aperiodic activity is causally involved in visual WM. In a randomized, double-blind, sham-controlled, crossover neurostimulation design, we showed that stimulating aperiodic activity with a steeper-than-average exponent resulted in improved visual WM precision, while post-stimulation resting-state activity was not affected by tRAS. Previous work using a similar task found increased aperiodic exponents during the delay period of a WM task, which could produce a confound to the typically reported increase in frontal-midline theta power (*23*). Thus, the delivery of steep-tRAS to improve WM precision is consistent with a role of the aperiodic exponent in the maintenance of WM contents.

While steep-tRAS improved visual WM precision as hypothesized, the relationship between trait-level aperiodic activity, the dynamics of the aperiodic exponent during a WM task, and the mechanism for how tRAS changes neural activity are complex. Multiple previous studies found that aperiodic activity was dynamically steepened during the delay period of delayed-match-to-sample or delayed-recall WM tasks (*23, 29*). However, other task factors produce a different pattern of aperiodic activity. For example, in a prospectively-cued WM tasks with a brief encoding and retention period the aperiodic exponent was decreased as a function of WM load (*17*). Another study investigating the dynamics of the aperiodic activity during memory encoding found that the exponent exhibits a canonical, approximately 1 second, decrease and return to baseline during stimulus presentation (*30*). With a 1-second encoding period in the task presented here, the delay period showed an increased aperiodic exponent. While our previous study showed a decrease in aperiodic exponent with only a 250 ms encoding period and 1300 ms delay period. This decrease could be caused by stimulus-related processes extending into the delay period (*17*). Nonetheless, we did not observe a load-dependent change in the aperiodic exponent which suggests that aperiodic signal changes may reflect a state-change that supports WM but is not the mechanism of WM per se. Consistent with this interpretation, steep-tRAS increased WM precision irrespective of the WM load.

Despite evidence that transcranial random noise stimulation (tRNS) can enhance learning and perceptual processing, there is limited evidence that conventional high-frequency tRNS reliably improves WM capacity or performance. Early work applying tRNS over the dorsolateral prefrontal cortex found no significant enhancement of WM, with performance improvements instead observed following tDCS (*31*). Similarly, pairing tRNS with repeated WM training did not increase either the magnitude of training-related improvement or transfer to untrained WM measures (*32*). Although Murphy et al. (2020)(*33*) reported improved Sternberg WM performance, their stimulation protocol combined high-frequency tRNS with a direct-current offset, making it difficult to attribute the behavioral effect specifically to random-noise stimulation. More recently, Tokikuni et al. (2024)(*34*) found that high-frequency tRNS without a DC offset improved dual n-back performance during stimulation; however, the dual n-back strongly engages executive updating and divided attention and may therefore differ from measures that more directly index WM capacity. Taken together, the existing literature does not provide consistent evidence that traditional high-frequency tRNS produces reliable or durable improvements in WM precision. Against this background, the present finding that steep-tRAS, with a prominent low-frequency component enhances WM performance whereas flat-tRAS, which more closely resembled tRNS, did not suggest that the cognitive effects of random-noise stimulation may depend critically on its spectral composition rather than reflecting a general benefit of adding neural noise.

We set out to investigate whether the aperiodic exponent was causally related to WM precision given the potential found that changes in aperiodic exponent produce for interpreting oscillatory effect. While theta power is known to subserve WM performance, the specific relationship with WM load which was observed in early studies has had mixed success in replication (*7, 8, 35*). Here, we found no increase in aperiodic-adjusted theta power as a function of WM load, nor in aperiodic exponent. Thus, theta activity in lateral frontal-parietal control networks may reflect additional control signals in the manipulation of WM content (*36*), output-gating (*37*), or conjunctive representations with how WM contents will be utilized (*38*).

While transcranial alternating current stimulation (tACS) is hypothesized to act on oscillatory activity via neural entrainment (*39*), tRNS is thought to alter neural excitability through stochastic resonance. The principle of stochastic resonance states that an optimum level of noise can be introduced to a system to increase its chances of detecting a subthreshold signal. At one extreme, too little noise and the signal of interest still does not surpass a detection threshold; at the other, too much noise overwhelms the system where noise itself surpasses a detection threshold (*40*). In fact, tRNS intensity follows an inverted U-shape pattern in terms of its efficacy (*41, 42*), and only increases detection of visual stimuli that were subthreshold, without modulating detection of suprathreshold stimuli (*43*). Recently, we posited that aperiodic neural activity might have a similar function in modulating neural activity (*21*). Given that aperiodic activity likely reflects the transmembrane currents integrating within a region, when the postsynaptic drive is dominated by longer timescale inputs – reflected by steeper spectra – the correlation structure increases, facilitating the integration of information. In contrast, when the postsynaptic drive is dominated by shorter timescale inputs – reflected by flatter spectra – the system is more driven by rapid inputs and is less effective at temporal integration. In terms of our current results, by steepening the spectra through tRAS, we may be pushing the underlying cortex into a state more amenable to integrating and sustaining information over longer timescales.

If stochastic resonance is the mechanism of action for tRAS, then the failure to produce a lasting impact on the aperiodic exponent is not unreasonable. By driving endogenous patterns of activity in an aperiodic-slope manner, for example low-frequency network-scale activity is facilitated by steep-tRAS and local modality specific processing is facilitated by flat-tRAS, there may not be a systematic change in the aperiodic signal following tRAS. In other words, the aperiodic signal is an emergent property of the brain, and as such cannot be directly entrained by an external driver. Instead, tRAS is mimicking a brain state that supports certain types of endogenous activity.

A few methodological limitations of our study must be addressed. One limitation is the intermixed trial design with each instance of tRAS to be delivered at a short duration of 5-6 minutes (although cumulative dose was 20-24 minutes per tRAS type) compared to typical protocols that deliver stimulation uninterrupted for 20-60 minutes (*44*). The rationale for this decision was that many previous studies have found a significant increase in WM performance using the selected, or similar, montage targeting theta oscillations (*15, 45, 46*). However, the benefit of using longer stimulation durations is that it increases the change to induce sustainable neuroplastic effects (*47*), which could explain why we were unable to find post-stimulation effects of the aperiodic exponent, as was found recently in a study using custom made waveforms mimicking aperiodic activity but with a sustained superimposed oscillatory rhythm (*48*). Lastly, while some studies attempt to regress out the tACS waveform to analyze online EEG effects, we avoided this method because the aperiodic waveforms are much harder to correct for than the simple sine waves used in tACS (*49, 50*).

Another limitation is that we used a fixed stimulation protocol instead of an individualized one. For instance, with tACS the stimulation center frequency can be individualized by mapping it to the center frequency of the individuals for the targeted neural oscillation (*51, 52*). Even though most participants were within the aperiodic exponent parameters for steep-and flat-tRAS (supplementary methods and Fig. S3), an individualized approach for tRAS could be beneficial (*48, 53*). Furthermore, flat-tRAS was somewhat comparable to tRNS and had no influence on behavior. Typically, tRNS protocols utilize a higher frequency ranges in their electrical waveform (100-700 Hz) (*54*), whereas we chose a frequency range within canonical EEG range (1-50 Hz). These discrepancies raise the possibility that stimulating with an exponent more aligned to endogenous aperiodic activity might have a stronger effect, similarly to what is seen in tACS (*55*).

Despite these complexities, we believe that our novel version of noninvasive transcranial electrical stimulation specifically designed to mimic aperiodic activity, tRAS, offers an innovative approach for causally studying the role of the aperiodic signal on human perception and cognition. While we did not find any sustained effects on aperiodic activity after the stimulation ended, we observed significant improvements in visual WM precision during steeper-exponent tRAS. Behavioral improvements in WM performance are generally rarer in the broader neurostimulation literature – it is easier to disrupt cognitive functioning than it is to enhance it. We are thus encouraged by the future possibilities for individualized, aperiodic neurostimulation for studying the role of aperiodic activity in cognition, as well as in the potential for causal manipulations of aperiodic activity in improving human cognitive functions. While we focus here on healthy younger adults, there are clear opportunities to study tRAS in the context of aging and age-related cognitive decline, as well as in disease-related disruptions to memory.

## Acknowledgments

We thank Research Assistant Shyam Dhulashia for collecting the data at Florida State University, and Research Assistant Gabriela Freedland for helping analyze the data at the University of California San Diego. We further want to thank Andrew Bender, Dillan Cellier, Ryan Hammonds, Blanca Martin-Burgos, Michael Preston, Eena Kosik-Rose, Sydney Smith and Christian Cazares for their advice and feedback on the manuscript.

## Author contributions

Conceptualization: QvE, JR, BV

Neurostimulation development: QvE, JR, BV

Data collection: QvE, JR

Data analysis: QvE, BV

Visualization: QvE, JR, BV

Funding acquisition: BV

Supervision: JR, BV

Writing – original draft: QvE, JR, BV

Writing – review & editing: QvE, JR, BV

## Data, code, and materials availability

All data are available in the main text or the supplementary materials. Data collected for this study can be downloaded from OSF (6qnfw). Custom scripts used in this study can be downloaded from github.com/voytekresearch/tras_wm

