## Supplementary materials for "Transcranial Random Aperiodic Stimulation Improves Working Memory Precision"

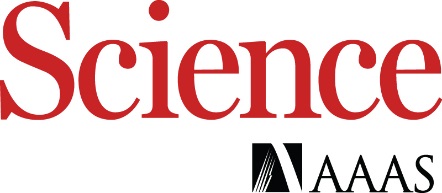

Supplementary Materials for

Transcranial Random Aperiodic Stimulation Improves Working Memory Precision

**Authors:** Quirine van Engen^1^*, Justin Riddle^2,3^†, Bradley Voytek^1,4-6^†

**The PDF file includes:**

Materials and Methods

Supplementary Text

Figs. S1 to S5

Table S1

References

Materials and Methods

**Participants**

Participants were recruited from the Tallahassee area and provided written informed consent upon enrollment in the study. A total of 35 participants enrolled in session 1. Our sample population had a mean age was 21土2.9, from which 27 were women, 7 men, and 1 identified as nonbinary. The majority of participants were right-handed, 33, and the other 2 were left-handed. Four participants were removed from further analysis because two failed to follow instructions, and another two only finished three or four out of the eight blocks. Two EEG recordings were corrupted and were excluded from further analysis, which resulted in a total of 31 participants for the behavioral analysis in session 1, and 29 for the EEG analysis.

Thirty participants from the 35 returned for session 2. In session 2, two were removed due to failing to follow instructions, and one for terminating the experiment prematurely. Two more participants were excluded from further analysis due to excessive noise in their EEG recordings. Thus, 27 participants were included in the behavioral analysis of session 2, and 26 for the EEG analysis.

*Inclusion/exclusion criteria*

All participants were between the ages of 18 and 35, had normal or corrected-to-normal vision, and were not color-blind. Participants were excluded if they were currently under treatment for ADHD/ADD, taking benzodiazepines, had a neurological disorder or those who had prior brain surgery, a brain device/implant (including cochlear implants and aneurysm clips), or any history of traumatic brain injury. Furthermore, females who were pregnant or breastfeeding were also excluded.

**Task design: continuous-report working memory task**

The behavioral task was programmed in MATLAB 2023A using Psychtoolbox version 3 (*1*–*3*). Participants performed a delayed continuous-report working memory task (Figure 1F). A baseline period of 2 sec was presented with a fixation cross in the middle of the screen. Then a 1 sec encoding screen appeared with a memory array showing three to seven colored squares. Participants were instructed to remember the color and position of each square during a delay of 2 sec. After this delay period, all locations were indicated with empty squares from which one of them was highlighted. The participant had to indicate the color of the highlighted square using a joystick moving a bar around on a color wheel within a 2 sec response window. Participants were instructed to be as fast and accurate as possible. The inter-trial-interval was jittered between 2 and 3 sec. From each trial, we collected the response time and the WM precision (error between the target and reported color).

There were 14 possible spatial locations for the WM items to appear, equally spaced around the color circle shown during the prompt screen. Nine possible colors were selected per block with an equal distance of 40 degrees from each other, with a 0 to 4 degrees random jitter added. All colors were matched for saturation and brightness.

**Procedures**

Participants were recruited to complete two days of recordings (Figure 1E). The first session served as a baseline session in which we assessed their individual WM capacity to titrate the task difficulty during the second session. Each participant received three stimulation types while performing the WM task during session 2: steep-tRAS, flat-tRAS, or sham. The sequence at which tRAS was administered was double-blind, randomized, and counterbalanced across participants. At the start of each block, steep-tRAS and flat-tRAS ramped up for 15 sec when the task began, continued to stay on, and ramped down for 15 sec after all trials within that block were completed. The sham condition included the ramp up and down protocol, but the stimulation stayed on for 30 sec instead of 5 min. This ramping protocol was repeated at the end of each block. We picked randomly from the steep-tRAS or flat-tRAS waveforms for this sham procedure. The shame protocol reproduced the perceived sensations on the skin (tingling, fade) induced by the active conditions and with the acclimation of neurons in the scalp to tonic stimulus, and thus, created an effective placebo condition (*4*, *5*). Note that with the interleaved block design we were limited in our ability to assess blinding. However, the inclusion of an active control condition in the form of steep-tRAS versus flat-tRAS precludes the possibility that our effects were driven by the placebo effect.

During session 1, the task conditions consisted of a WM load of either 3, 4, 5 or 6 items. The order of loads was randomized and intermixed within block. There were 8 blocks and 40 trials per block, resulting in a total of 320 trials in total, and 80 trials per load. Each block took 5 min to complete the 40 trials and was followed by a 2 min resting period where participants gazed at a fixation cross and let their mind wander. As mentioned before, individual WM capacity was determined by their performance in session 1. Accuracy was calculated as the percentage of trials (excluding misses) with a WM precision under 30 degrees for a load condition. If load-4 accuracy was less than 50% or if load-5 accuracy was less than 60%, participants were assigned the low-load group (3 and 5 items). Otherwise, if load-6 accuracy was less than 70%, participants were placed in the medium-load group (4 and 6 items). Otherwise, participants were placed in the high-load group (5 and 7 items). In addition, participants with a performance near chance levels or with an excess of non-responses (25%) were excluded from the study and did not complete the second session.

During session 2, the task conditions consisted of either 3 versus 5, 4 versus 6, or 5 versus 7 items depending on the titrated WM load from the baseline session. The low and high loads were randomly intermixed within each block. This session consisted of 12 blocks and 40 trials per block. There was a total of 480 trials with 240 trials per individualized low or high WM load. Each block in session 2 took about 5 min to complete and either steep-tRAS, flat-tRAS or sham was administered for each block, followed by a similar 2 min resting period.

**EEG acquisition**

EEG was recorded during both sessions with a 96 active channel Easycap and the ActiChAmpPlus from BrainVision (actiCHamp Plus, Brain Products GmbH, Gilching, Germany). Electrode Fpz was used as ground, and Pz as online reference. Data were recorded with a sampling frequency of 1000 Hz. HOG and VOG channels were included in the setup to account for eye movements and blinks during pre-processing. Additionally, a photodiode channel was used for synchronization of the EEG to the LCD monitor. Participants' chins were placed in a chinrest to maintain a fixed distance from the monitor.

**Electrical stimulation**

*Electrical stimulator details*

We used the neuroConn DC-Stimulator MultiChannel from the Neurocare group (Neuroconn Ltd., Ilmenau, Germany) during session 2. A custom-built Y-cable was used for the return electrode. We used two 1 mA current generators.

*Stimulation montage*

The goal of the stimulation montage was to target the left lateral frontal-parietal network (Figure 1D), due to successful modulations in WM using theta tACS. The target stimulation pads were placed over F3 and P3 (according to the international 10-20 system), and sized 4.5 by 4.5 cm. The return pad was 5 by 7 cm and placed over Cz (central midline). This montage was selected based on previous work using this or a similar montage that found an improvement in WM performance during theta-frequency tACS (*6*, *7*).

*Transcranial Random Aperiodic Stimulation (tRAS) waveform creation*

We used a fixed stimulation protocol for each participant. The steep-tRAS condition was designed to mimic a steeper power spectrum than the average endogenous activity, and vice versa for flat-tRAS. Based on a re-analysis of a previous EEG experiment using a visual WM task (*8*, *9*), the average aperiodic exponent during the delay period in one experimental cohort was 1.14±0.36 μV^2^/Hz during the delay period and 1.17±0.35 μV^2^/Hz for the cue period. For a second independent experimental cohort, the exponent during the delay period was 1.08±0.36 μV^2^/Hz and 1.13±0.30 μV^2^/Hz for the cue period, all measured on frontal midline electrodes (F3, F4, and Fz). To make sure there was an equal distance from endogenous aperiodic activity and the two stimulation conditions, flat-tRAS was created with an exponent of 0.8 μV^2^/Hz, whereas steep-tRAS had an exponent of 1.6 μV^2^/Hz.

Based on the data from the resting periods of the baseline session, we checked whether the current participants’ endogenous aperiodic exponents were within the predetermined settings described above. Most participants were well within the 0.8 and 1.6 μV^2^/Hz tRAS exponent settings (Fig. S3). Only four participants had an endogenous exponent that was higher than our steep-tRAS parameter, and only two other participants had an endogenous exponent that was lower than the flat-tRAS parameter.

The tRAS waveforms were created in Python using toolboxes numpy and scipy. Ten electrical stimulation protocols were created by applying the inverse FFT of the aforementioned exponents with random phases for 1 to 50 Hz with 0.01 frequency spacing, resulting in a sampling frequency of 10,000 Hz. This signal was 6 min long to ensure stimulation was on during the entire task block. The average stimulation value was 1 mA (2 mA peak-to-peak) without a DC-offset, delivered through two channels for a total current of 2 mA (4 mA peak-to-peak). We deliberately chose a frequency range between 1-50 Hz since this is the frequency range typically analyzed with scalp-EEG. Notably, this deviates from most tRNS protocols that deliver current with power in the high frequency range (starting at 100 Hz, and up to 700 Hz) (*10*).

One previous study used “pink-noise” electrical stimulation, where pink noise followed a 1/f distribution that was designed to resemble a steeper-than- endogenous neural aperiodic activity. They found pink-noise stimulation inhibited food-cravings (*11*). Thus, these researchers demonstrated that stimulating a steeper spectral distribution produces systematic changes in behavior, without causing adverse side effects.

**Behavioral analysis**

The Target Confusability Competition model (TCC) (*12*) was used to analyze the behavioral data. This model was appropriate for analyzing continuous visual WM responses since it uses the distance between the colors in combination with signal detection theory to calculate the memory precision (d′). Higher memory precision was indicated with a higher d′ value. The input for this model was the WM precision data for each condition, load in session 1, and load and stimulation type in session 2 (Figure 1G).

We observed that participants had significantly more missed responses (not answered within the 2 sec window) in the higher loads than the lower loads in session 1 (β=2.87, p=2.72*10-6, 95%CI = [1.62, 4.15]), but this effect was not present in the stimulation session for WM load (F(1, 26)=3.06, p=0.092, η_p_^2^=0.11). Stimulation type did not change the number of missing responses (F(2, 52)=0.17, p=0.84, η_p_^2^=0.006). The ratio of missing responses based on the total number of trials recorded per participant and conditions was neither significant for the WM load (F(1, 26)=3.38, p=0.078, η_p_^2^=0.12), nor for the stimulation type (F(2, 52)=0.33, p=0.71, η_p_^2^=0.013). Nevertheless, missing responses are problematic for accurate memory precision estimation by the TCC model and can skew results between the conditions of our procedure. To account for missing responses in the TCC model, we assumed that a missing response was equivalent to a guess. To model this, we inserted random values drawn from a uniform distribution between -180 and +180 error degrees for 1000 simulations. For each simulation, the TCC model was applied as previously described, on each condition within each participant. Then, the averages were taken to get a more accurate d′ estimate before statistical analysis.

**EEG preprocessing**

*Session 1 – baseline*

A FIR high-pass filter of 0.5 Hz was applied before down sampling the data from 1000 Hz to 500 Hz. After this, noisy electrodes were identified by visual inspection of the power spectra and the time series per electrode. Next, we re-referenced data to the common average before applying fast ICA. The number of components was determined as one less than the number of non-interpolated electrodes and as such varied by participant based on the number of noisy electrodes identified. The average number of components in session 1 was 87 土 6. ICA components related to eye movements or blinks, and other non-neural artifacts, were removed. Lastly, noisy electrodes were interpolated using the spherical spline method. After these preprocessing steps, data were epoched using the photodiode channel to extract the exact timing of the task and the 2 min resting-state period. The task-based data was time-locked to the onset of the memory array and epoched 3 sec before, and 4 sec after to include the pre-stimulus period (2 sec), the encoding (1 sec) and delay (2 sec) period. The resting period was also extracted as 2 min periods.

*Session 2 - stimulation*

Similarly to session 1, a FIR high-pass filter of 0.5 was applied followed by down sampling to 500 Hz. Since EEG recordings are heavily disrupted by the ongoing tRAS stimulation, data were epoched to the resting-state periods when stimulation was off. Then, noisy electrodes were identified visually by looking at the raw time series and power spectra. Noisy electrodes were interpolated using the spherical spline method before re-referencing to the common average. Fast ICA was applied to remove eye movements and blinks, or other non-neural artifacts. In session 2, the average number of ICA components was 63 土 9. Session 2 ICA used fewer components than session 1, because electrodes near the stimulation pads were not connected to avoid bridging.

**EEG analysis**

*Computing power spectral density (PSD)*

We obtained PSDs by using the FFT in MNE with a 2 sec Hamming window without overlap for both the pre-stimulus and delay period for the task data from session 1. The time-resolved PSDs from the 2 min resting-state period from session 1 and 2 were obtained by applying the short-time FFT from SciPy signal using a 5 sec window with 4.5 sec of overlap.

*Spectral parametrization*

To separate contributions from oscillatory and aperiodic dynamics, we applied spectral parameterization, using specparam (*13*). The model was fit between 2 and 35 Hz to exclude common muscle artifacts in EEG. The maximum number of peaks fit was 12. These peaks had to be between 1 and 8 Hz in width, a minimum peak height of 0.05, and a soft peak threshold of twice the standard deviation from the mean. Data was fit without a knee, given visual inspection of single trial PSDs did not show a clearly defined bend. Theta oscillations were defined between 4-7 Hz, and alpha between 8-12 Hz. All these settings were kept constant over conditions and participants.

Aperiodic activity was captured by the exponent and offset. Oscillatory power was calculated as the area under the curve (AUC) between the fitted aperiodic activity and the remaining PSD values within a frequency band of interest, theta or alpha (Figure 1H). Furthermore, total band power was also calculated as the AUC, but without correcting for aperiodic activity for easier comparisons with the existing literature. In session 1, these outputs were collected from each trial, time window, and electrode. Models were fitted on the single-trial generated PSDs. Baselining was performed by subtracting the output from the pre-stimulus period from the delay period windows. For resting data, the same outputs were collected, but for each electrode and time window within the 2 min resting period. Here, PSDs were averaged per time window and electrode within a condition before model fitting.

Model fitness was evaluated using the goodness-of-fit metric (R^2^). For the single-trial fitting in session 1, windows with a R^2^ below twice the standard deviation from the mean were excluded from further analysis. For a trial and electrode combination to remain included, both the pre-stimulus and the delay window fits had to be above the aforementioned threshold. For the resting state, only time windows of electrodes below three times the standard deviation from the mean were removed from further analysis. Spectral parameterized time series from rest were further cleaned by removing the first 5 and last 5 sec of the series to remove edge artifacts.

Electrode clusters were defined by visually inspecting the topoplots or determined by the placement of the stimulation pads in session 2. A frontal-midline cluster was chosen to include Fz, AFF1h, AFF2h, FFC1h, FFC2h, FC1, and FC2 to investigate theta oscillatory activity. The aperiodic exponent was investigated on a left frontal-central cluster defined by F3, FC1, FC3, C1, C3, C5. Clusters centered on the stimulation pads were defined as a left frontal cluster (F3, F1, F4, FFC5h, AF3), and a left parietal cluster (P3, P1, P5, CPP3h, PO3), because the target stimulation pads were placed on F3, and P3.

**Statistical analysis**

All statistical analyses were performed in Python, using the pingouin package, except for the Linear Mixed Models (LMM) which were performed in R using lmerTest.

In session 1, the independent variable was the WM load, either 3, 4, 5 or 6 items. The dependent variables were the working memory precision (d′) from the TCC model, the aperiodic exponent, and aperiodic-corrected theta power. Given the order in the WM load, a Linear Mixed Model was applied to test for significant differences between the loads.

We had two levels of independent variables in session 2, the load – either low or high – and the stimulation types – steep-tRAS, flat-tRAS, or sham. The dependent variable was the WM precision (d′) from the TCC model. A two-way repeated measures ANOVA was used to determine whether stimulation type and individualized WM load influenced WM precision. If data were not spherical, a Greenhouse-Geisser adjustment was applied on the calculated p-values. If the effects were significant, a pairwise post hoc test was performed with a Holm-Bonferroni correction for multiple comparisons.

To further assess the effect of stimulation, we calculated the difference in d′ between the steep-tRAS and flat-tRAS conditions (d′ slope) per participant and load. A positive slope in d′ indicated improved WM precision in the steep-tRAS condition, whereas a negative slope indicated an improvement due to flat-tRAS.

Any correlations were calculated using Spearman’s rank correlations. In session 1, we correlated either the baselined aperiodic exponent or the baselined aperiodic-corrected theta power to the working memory precision per load and added a Holm-Bonferroni correction for multiple comparisons. This was to test for associations between individual task-related EEG effects within loads over participants. Next, for assessing whether resting aperiodic activity was correlated to WM precision, we first averaged d′ over loads per participant and correlated the resulting value with their exponent during rest in the same session, session 1. No multiple comparison adjustments were applied. Last, to further assess if aperiodic activity at rest influenced stimulation efficacy, we correlated the aperiodic exponent at rest during session 1, to the WM precision difference from flat- to steep-tRAS. Two Spearman correlations were applied per individualized load, with a Holm-Bonferroni correction.

**Software**

The behavioral experiment was coded in MATLAB (2023b) using Psychtoolbox (*1*–*3*). The TCC model was performed using MemToolBox (*12*, *14*) in MATLAB. The LLMs were performed in R (v2023.06.0+421) with Lmetest (v3.1.3). All other analysis, including the tRAS waveform creation were performed in Python (v3.8.5). The MNE (v1.6.1) toolbox (*15*) was used for pre-processing EEG data. Data were analyzed using Pandas (v2.0.3) (*16*), NeuroDSP (v2.2.1) (*17*), and Specparam (v2.0.0rc1) (*13*). Statistics, except LMMs, were executed with the Pingouin (v0.5.4) package (*18*). Stimulation waveforms were created using numpy (1.24.4) and Scipy (v1.10.1) (*19*). All data was visualized with Matplotlib (v3.7.5) (*20*) and Seaborn (v0.13.2) (*21*).

Supplementary Text

**Adverse side-effect analysis**

In the event of a participant reporting a side effect (4 out of 0-4 on a Likert scale), a structured adverse events interview was conducted. Four out of thirty participants reported an adverse event that was deemed to be mild. Two reported sleepiness, one reported sleepiness and a burning sensation, and the last one reporting tingling and flickering. An overview of all answers can be found in Table S1.

**tRAS exponent parameter check**

Given that we a priori picked the exponent parameters for steep- and flat-tRAS based on a re-analysis of a WM study (Adam et al., 2018; van Engen et al., 2026), we checked whether the measured aperiodic exponent values were comparable in the current study. For this, we calculated the aperiodic exponent during session 1 for the electrodes that would be placed on top of the stimulation pads in session 2: the left frontal and left parietal clusters. We found an average exponent of 1.33 μV^2^/Hz, which was steeper than the re-analysis, which found 1.13 μV^2^/Hz. Note that different electrode clusters were compared, which could have contributed to this discrepancy. Nevertheless, most participants had a measured aperiodic exponent that fell within the chosen tRAS parameters of 0.8 μV^2^/Hz and 1.6 μV^2^/Hz (Fig. S3). Only five out of twenty-nine participants had aperiodic activity with a higher exponent than the steep-tRAS condition, and two other participants had an exponent lower than the flat-tRAS condition.

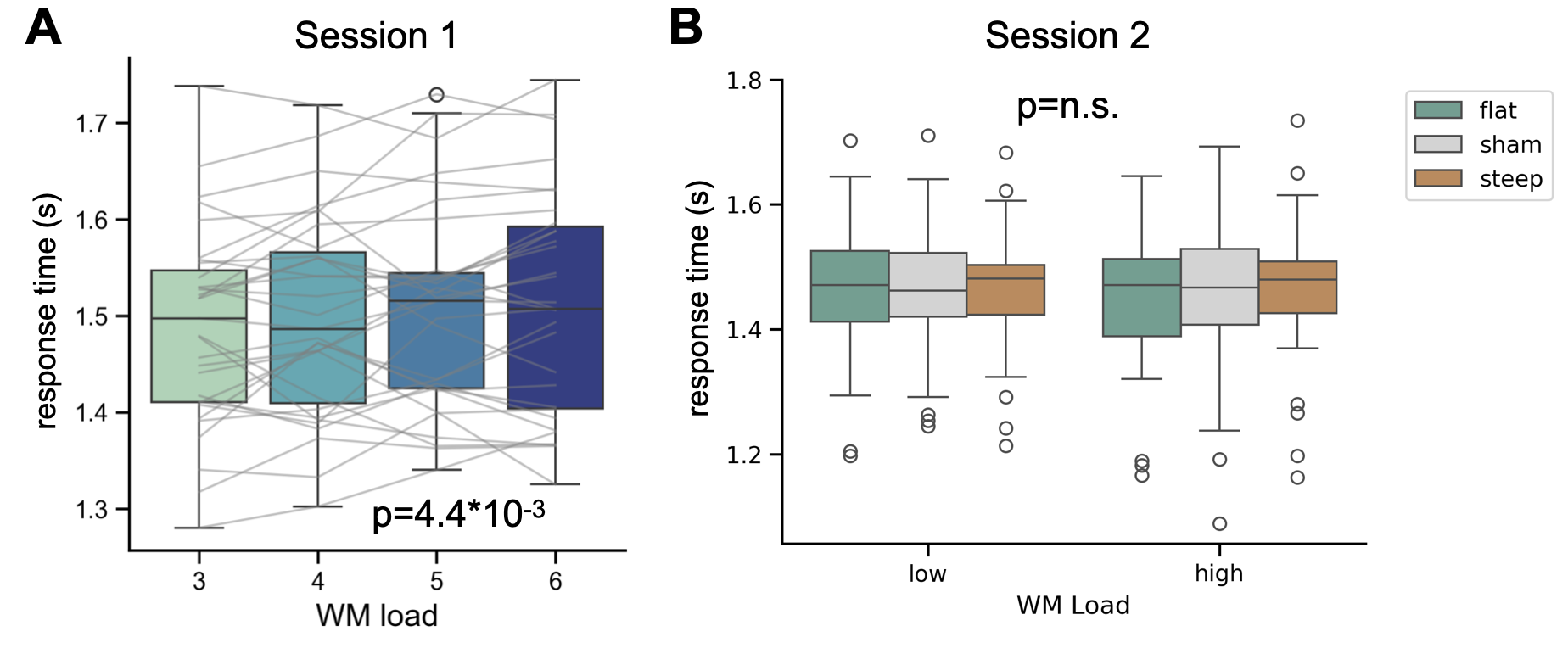

Fig. S1. Response times increased with higher WM loads during session 1, but not during session 2. No effect of stimulation type was found on response time in session 2 either. A linear mixed model was used for session 1 (β = 0.0087, p = 4.4*10-3, 95%CI = [0.003, 0.015]), and a repeated measures ANOVA on the session 2 data. There was no main effect of WM load on response time (F(1,26) = 2.4, p = 0.13), nor a main effect of stimulation type (F < 1).

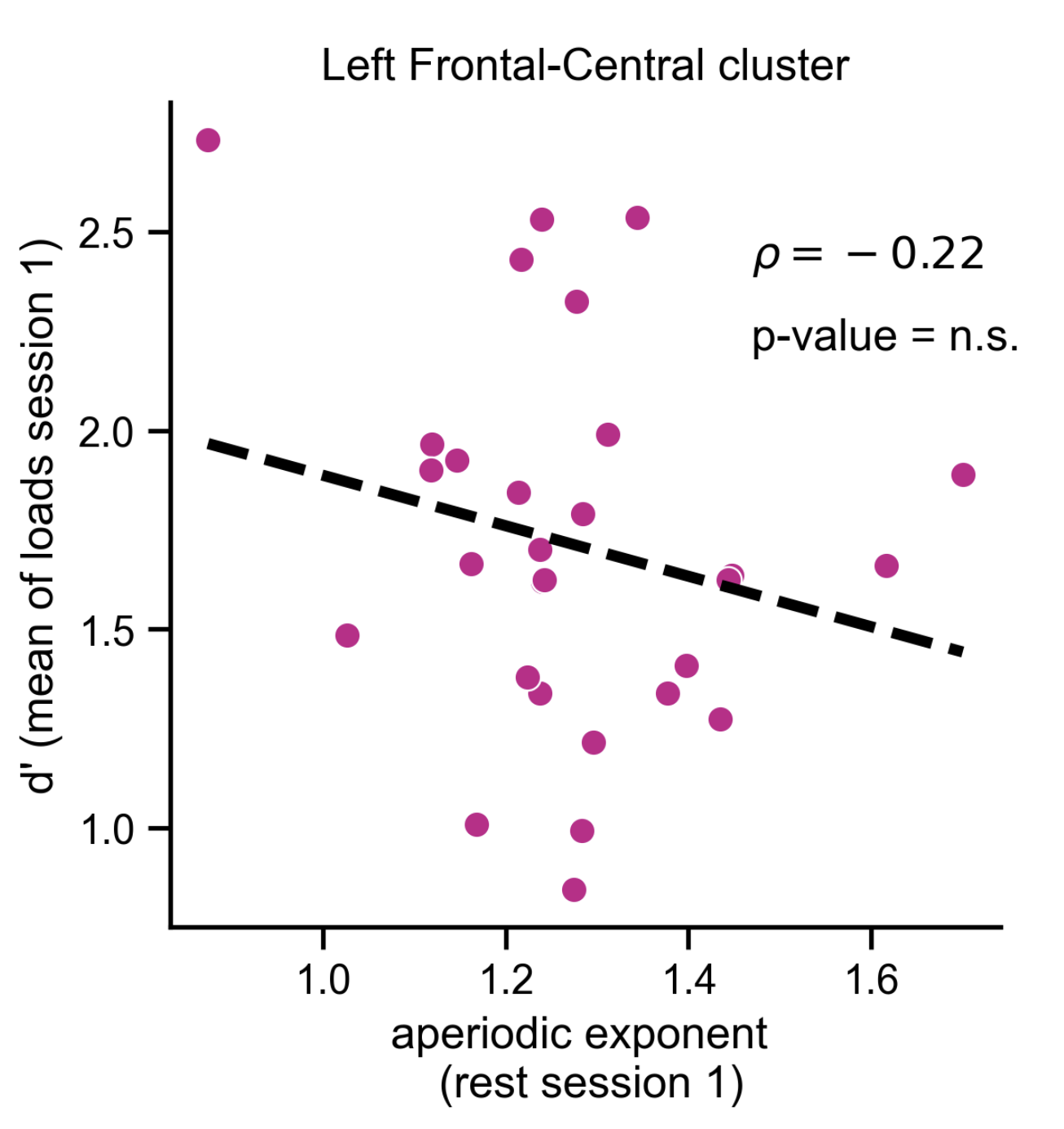

Fig. S2. Aperiodic activity at rest in session 1 did not predict WM precision. Each dot represents a participant's aperiodic exponent at rest (in between blocks) in session 1, and their average WM precision over the four loads in session 1. The correlation analysis revealed a negative relationship, but the association was not significant. Thus, resting aperiodic activity was not associated with overall WM precision.

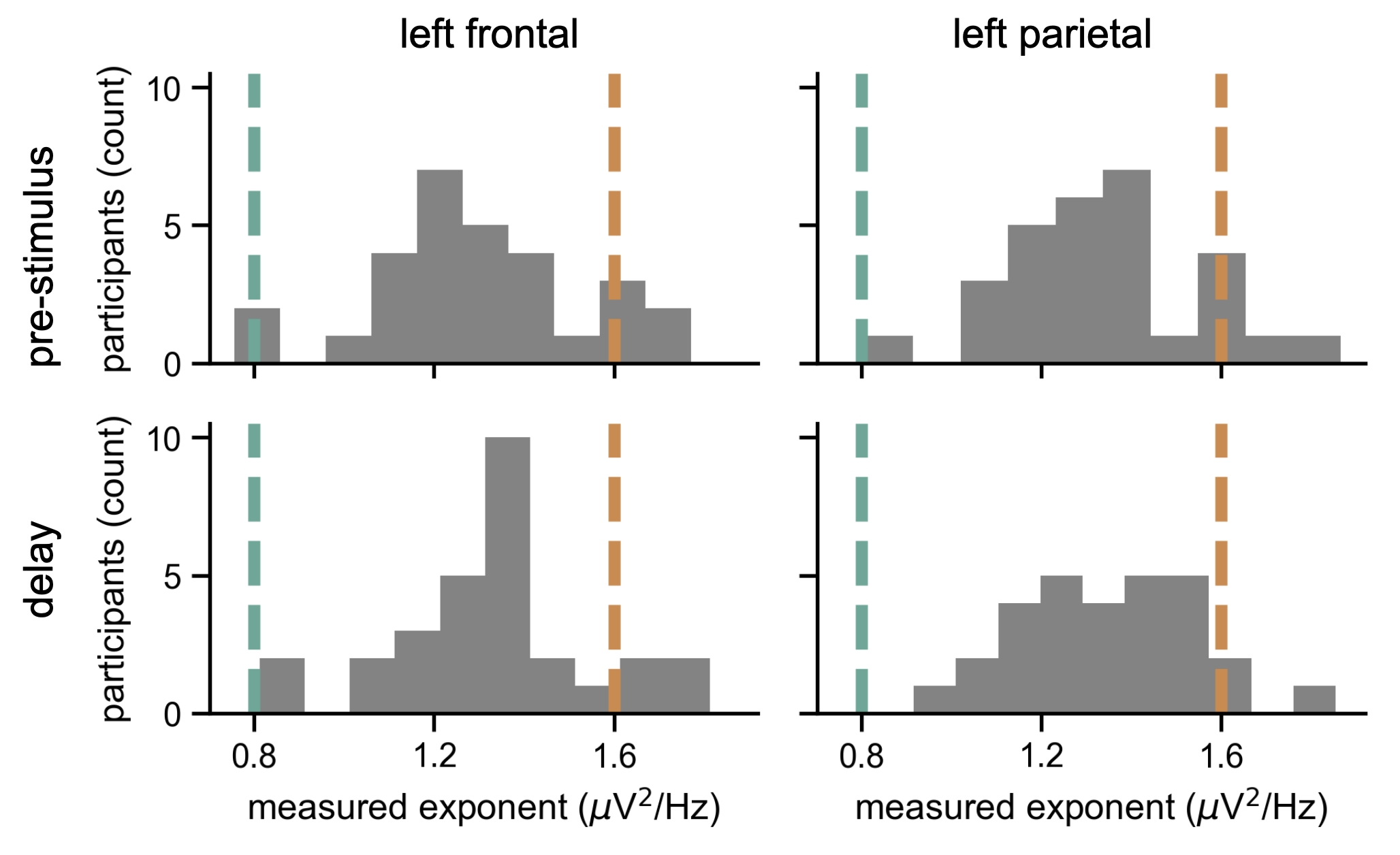

Fig. S3. Measured aperiodic exponent was mostly between the fixed tRAS exponent parameters. We chose our tRAS exponent parameters a priori based on the re-analysis of another WM study (Adam et al., 2018; van Engen et al., 2026). The average aperiodic exponents found over the frontal-midline in that study was around 1.13 μV^2^/Hz. The exponent for flat-tRAS was set at 0.8 μV^2^/Hz, and steep-tRAS was 1.6 μV^2^/Hz to ensure equal distance between the average measured aperiodic exponent, and the stimulation parameters. In the current study, the average aperiodic exponent over left frontal and parietal regions, and the pre-stimulus and delay period was 1.33 μV^2^/Hz.

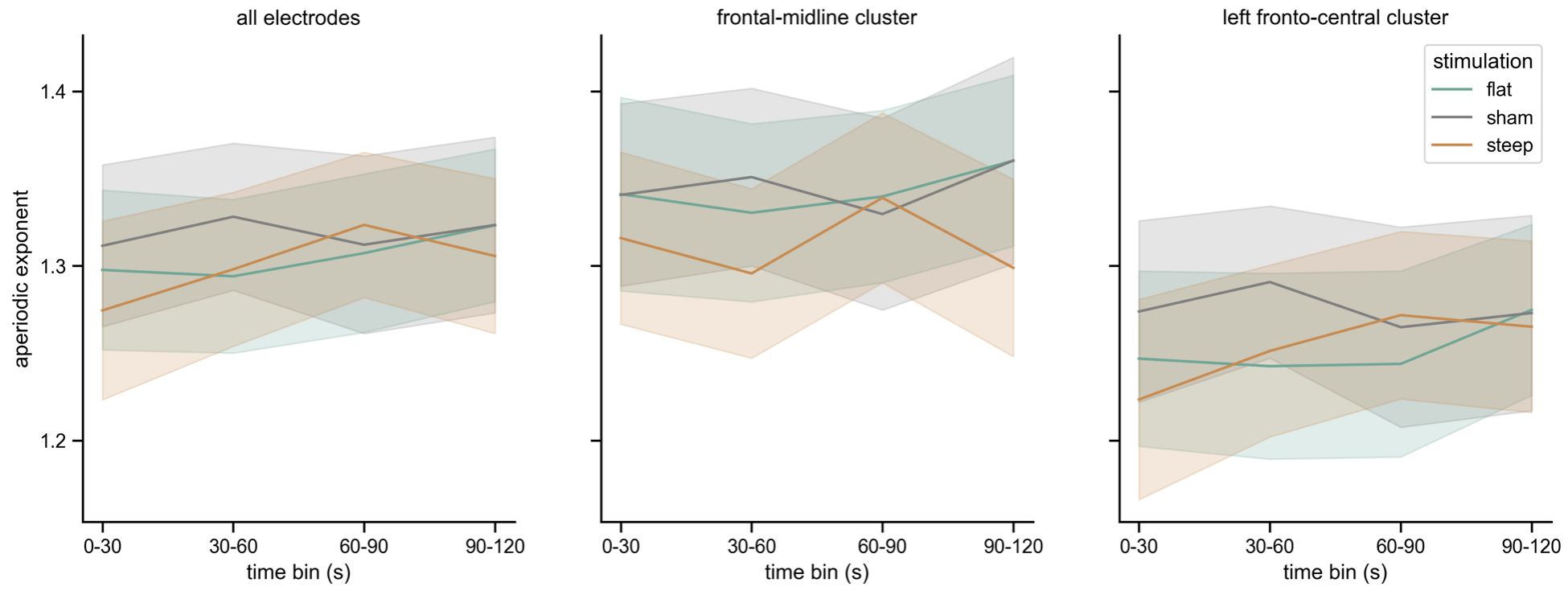

Fig. S4. Post-stimulation aperiodic exponent did not show a significant difference between stimulation types. The aperiodic exponent was investigated over four time bins during the post-stimulation resting period. No significant effect of stimulation type on aperiodic exponent was found.

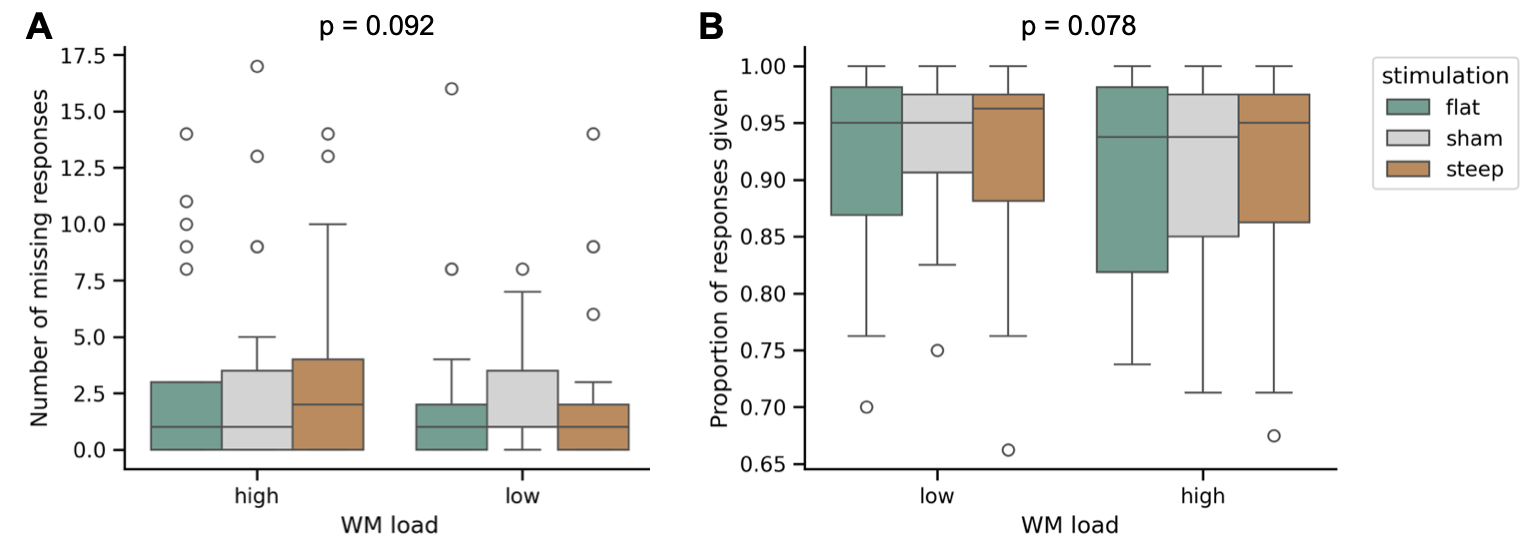

Fig S5. Response rate was affected by neither WM load nor stimulation type. (A) The number of missing responses was investigated for each condition. (B) Similarly, the proportion of responses provided was investigated. P-values in the figures represent the main effect of WM load.

Table S1. Side effects of tRAS. The number of participants that experienced each side effect and the severity on a 0-4 Likert scale for all 30 participants. 0 = absent.

| **Side effect / likert scale rating** | **0** | **1** | **2** | **3** | **4** |
| --- | --- | --- | --- | --- | --- |
| Headache | 21 | 7 | 1 | 1 | 0 |
| Neck pain | 22 | 4 | 3 | 1 | 0 |
| Scalp pain | 14 | 9 | 5 | 2 | 0 |
| Tingling | 4 | 7 | 13 | 5 | 1 |
| Itching | 17 | 7 | 5 | 1 | 0 |
| Ringing/buzzing noise | 24 | 4 | 2 | 0 | 0 |
| Burning | 10 | 11 | 5 | 3 | 1 |
| Sleepiness | 1 | 10 | 7 | 9 | 3 |
| Trouble concentrating | 3 | 8 | 12 | 7 | 0 |
| Improved mood | 25 | 0 | 4 | 1 | 0 |
| Worsening of mood | 25 | 2 | 4 | 1 | 0 |
| Dizziness | 23 | 6 | 1 | 0 | 0 |
| Flickering lights | 15 | 4 | 6 | 4 | 1 |
| Local redness | 20 | 9 | 1 | 0 | 0 |
